# Sex differences in human IgG1-mediated angiogenesis inhibition depend on Y chromosome-encoded DDX3Y

**DOI:** 10.1101/2025.03.25.645206

**Authors:** Dionne A. Argyle, Roshni Dholkawala, Ryan D. Makin, Felipe Pereira, Yosuke Nagasaka, Peirong Huang, Shao-bin Wang, Ivana Apicella, Abhinav Arneja, Chance John Luckey, Kenneth A. Walsh, Arthur P. Arnold, Jayakrishna Ambati, Bradley D. Gelfand

## Abstract

Prevalent diseases of angiogenesis such as age-related macular degeneration (AMD) exhibit sex-specific prevalence, with female sex being an independent risk factor for the advanced neovascular form of AMD. The basis for these sex differences is poorly understood. Here, we quantify the impact of sex on suppression of aberrant choroidal angiogenesis by human immunoglobulin 1 (hIgG1), which possesses class-wide, antigen-independent anti-angiogenic activity. Males exhibited significantly more robust hIgG1-induced chemotaxis inhibition in primary human and mouse macrophages and anti-angiogenic activity in mouse laser-induced choroidal neovascularization (CNV). Four core genotypes and XY* Turner Syndrome mouse models revealed the Y chromosome as a sex-biasing factor contributing to hIgG1 responses. Transcriptomic and RNAi screening identified DEAD-Box Helicase 3 Y-Linked (DDX3Y) as necessary for the robust inhibitory effects of hIgG1 in males. Male mice as well as human and mouse macrophages lacking DDX3Y exhibited blunted hIgG1 responses in CNV and macrophage chemotaxis, resembling those of females. These results unveil a novel mechanism through which sex chromosome complement differences can impact key processes involved in AMD, carrying implications for various sex-related disorders and conditions involving angiogenesis.

## Introduction

Age-related macular degeneration (AMD) is a leading cause of blindness with an estimated worldwide prevalence of 2.5%.^1^ Neovascular AMD is an advanced disease stage that accounts for 90% of legal blindness due to AMD.^2^ Neovascular AMD is caused primarily by choroidal neovascularization (CNV), a pathological process in which aberrant neovessels invade the outer retina, which is normally avascular. These fragile neovessels compromise vision by inducing edema, hemorrhage, and fibrosis.

Established risk factors for AMD include age, genetics, tobacco consumption, and female sex.^2^ Among European populations, females are at a significantly increased risk of developing neovascular AMD.^3–6^ However, the mechanistic basis for female predisposition to neovascular AMD remains unclear. Associations between AMD and menarche and menopause ages and duration, and hormone replacement therapy are variable and inconsistent (reviewed in ^7^). Mouse studies of experimental CNV report that sex does not affect CNV in relatively young animals, though middle-aged females develop greater pathology than males.^8–10^ Though some evidence suggests that estrogens may contribute,^8^ the potential role of sex chromosome complement has not been investigated previously.

Human immunoglobulin 1 (hIgG1) and its murine counterparts, IgG2a and IgG2c, inhibit choroidal neovascularization independently of antigen binding. This inhibition arises from the interaction between the hIgG1 Fc domain and macrophage-expressed Fc gamma receptor 1 (FcγR1), resulting in reduced VEGFR1 expression, blunted VEGF-A-driven macrophage chemotaxis, and reduced angiogenesis.^11^

Here we report robust sex differences in the inhibitory activity of hIgG1 for human and mouse macrophage chemotaxis and CNV in young mice, with hIgG1 eliciting substantially more angiogenesis inhibition in males compared to females. Using genetic models, we report sex chromosome complement as a contributing sex-biasing factor to this difference. We identified the Y chromosome-encoded gene DDX3Y as a critical mediator of hIgG1-induced angiogenesis inhibition using genetic and transcriptomic screens. These findings reveal a functional role for a Y chromosome-encoded gene as responsible for sex differences in angiogenesis regulation.

## Methods

### Mice

All animal experiments were approved by the University of Virginia’s Institutional Animal Care and Use Committee. Male and female mice wild-type (WT) C57BL/6J, *Fcgr1*^−/−^ (Jackson Strain No. 000664), B6D2F1/J (Strain No. 100006), and BALB/cJ (Strain No. 000651) were purchased from Jackson Laboratory. The four core genotypes source strain, B6.Cg-Tg (Sry) 2Ei Sry^dl1Rlb^/ArnoJ mice (Jackson Strain No. 010905), were previously described.^12,13^ Turner syndrome model XY* mice were previously described (MMRRC Strain 043694-UCD).^13,14^

### Cell Culture

Human male THP-1 cells were cultured in RPMI-1640 media (ThermoFisher No. 118754119) supplemented with 10% fetal bovine serum (FBS) and 1% penicillin-streptomycin. Human female HL-60 cells were cultured in IMDM media (ThermoFisher No. 12440061) supplemented with 20% FBS and 1% penicillin-streptomycin. RAW 264.7 and J774 cells were cultured in DMEM media (ThermoFisher No. 12430062) supplemented with 10% FBS and 1% penicillin-streptomycin. Primary bone marrow derived macrophages (BMDMs) were isolated from 4-week-old mice as previously described,^15^ and cultured in IMDM media with 10% FBS and 30% L929 conditioned supernatant, nonessential amino acids, sodium pyruvate, 2-mercaptoethanol, and antibiotics. All cells were maintained at 37°C in a 5% CO_2_ environment.

### Primary Human Macrophages

Buffy coats obtained from the venous blood of healthy donors were provided by A. Criss (University of Virginia) or from apheresis leukoreduction collars purchased from Crimson Core (Brigham and Women’s Hospital). Material used in this study were considered discarded material from other studies and exempt from human subjects oversight. Peripheral blood mononuclear cells (PBMCs) were isolated using Ficoll-Paque PLUS density gradient (Cytiva). To maximize the number of monocytes collected, RosetteSep® was added to the buffy coat for 20 minutes with incubation at room temperature. The sample was then diluted with PBS + 2% FBS (Stemcell) at the 2 times the initial volume then mixed. The diluted sample was added to the Ficoll® and spun for 20 minutes at 1,200 x g at room temperature with the brake off. Cells were washed with PBS + 2% FBS and collected. CD14+ cells were then selected for using the EasySep™ HLA Chimerism Buffy Coat CD14 Positive Selection Kit (Stemcell) following the manufacturer’s instructions. Cells were grown in DMEM/F12 media + 10% human serum + 1% penicillin-streptomycin + M-CSF for 5-7 days for macrophage differentiation and maintained at 37°C in a 5% CO_2_.

### Real-time PCR

Total RNA was purified from cells with either TRIzol reagent (Invitrogen) or RNeasy Micro Kit (Qiagen) according to the manufacturer’s recommendations and reverse transcribed with a QuantiTect Reverse Transcription kit (Qiagen). The RT products (cDNA) were amplified by real-time quantitative PCR (Applied Biosystems 7900 HT Fast Real-Time PCR System) with Power SYBR green Master Mix. Relative gene expression was determined by the 2^−ΔΔCt^ method, with Beta-Actin, 18S, or GAPDH used as an internal control.

### Macrophage Migration Assay

Primary mouse and human macrophages and differentiated THP-1, HL-60, RAW 264.7, and J774 cells were treated with trypsin, suspended in 2% BMDM medium and seeded onto the upper chamber of an 8 μm polycarbonate filter such that 5×10^3^ cells were seeded onto a 24-well or 2 × 10^4^ cells were seeded onto a 12-well format filter. Medium containing bevacizumab (0.1 mg/ml), hIgG1 purified from human myeloma plasma (0.1 mg/ml; Athens Research), or hIgG2 purified from human myeloma plasma (0.1 mg/ml; Athens Research) was placed in the lower chamber for 4 hours prior to the addition of recombinant mouse Vegf-a 164 (50 ng/ml; R&D Systems) or recombinant human VEGF-A 165 (50 ng/ml; R&D Systems). After allowing cell migration for 12-16 hours, cells were removed from the upper side of the membranes, and nuclei of migratory cells on the lower side of the membrane was stained with DAPI. The entire filter was imaged with a Nikon A1R epifluorescent microscope and the number of migratory cells were quantified using ImageJ.

### Laser photocoagulation-induced choroidal neovascularization (CNV)

Except where indicated, CNV was performed on mice up to 8 weeks old. As previously described,^16^ laser photocoagulation (532 nm, 180 mW, 100 ms, 75 μm) (OcuLight GL; IRIDEX Corp., Mountain View, CA, USA) was performed bilaterally (four spots per eye) on day 0 to induce CNV. 25 μg in 1 μL of bevacizumab or an equivalent volume of PBS were injected into the vitreous humor of mice using a 33-guage double-caliber needle once, immediately after laser injury. Where indicated, cell-permeable cholesterol-conjugated siRNA targeting Ddx3y or scrambled control was administered via intravitreous injection. 7 days after injury, mice were euthanized, and eyes were enucleated and fixed with 4% paraformaldehyde for 30 minutes at 4°C. Eyecups were incubated with fluorescein isothiocynate (FITC)-isolectin B4 (Vector Laboratories, Burlingame, CA, USA). CNV volume was visualized by scanning laser confocal microscopy (TCS, SP, Leica). Volumes were quantified using Image J software (http://imagej.nih.gov/ij/; provided in the public domain by the National Institutes of Health, Bethesda, MD, USA) as previously reported.^16^

### Gonadectomy surgeries

Orchiectomy, ovariectomy, and sham surgeries were performed by the Jackson Laboratory on 4-week-old male and female C57BL/6J mice. Mice were given a 28-day recovery period before CNV surgery was performed.

### siRNA transfection

siRNA transfection was performed using DharmaFECT 4 (Dharmacon) according to the manufacturer’s instructions. MISSION siRNA Universal Negative Control #1 (Sigma) and Control siRNA-A (Santa Cruz) were used as negative controls. Target siRNA for Ddx3y/DDX3Y, Uty/UTY, Kdm5d/KDM5D, and Rbm31y were manufactured by Dharmacon. Knockdown of targets was assessed by RT-PCR.

### Statistical Analysis

Statistical tests, analyses and graph generation were performed with GraphPad Prism 10 software. Outliers were assessed by Grubb’s test. Comparisons between two groups were performed with unpaired two tail t-tests for normally distributed datasets, or Mann-Whitney U tests for CNV volumes which are not normally distributed ^16^. Comparisons between more than two groups were performed using ANOVA with Šídák’s multiple comparisons test for multiple comparisons. P values < 0.05 were considered statistically significant.

## Results

### hIgG1 elicits a sex-specific anti-angiogenic effect

hIgG1 suppresses macrophage chemotaxis toward a VEGF-A gradient,^11^ which is an essential pro-angiogenic process.^17–19^ This hIgG1 activity is independent of antigen binding, but instead occurs due to monomeric hIgG1 binding its high affinity receptor FcγR1/CD64, which suppresses the expression of the VEGF-A receptor VEGFR1.^11^ Bevacizumab is a full-length humanized (94% human, 6% murine) hIgG1 antibody that neutralizes human VEGF-A but has essentially no affinity for Vegf-a from mice due to amino acid differences between mouse and human in the antibody-VEGF-A binding regions.^11,20,21^ Thus, in mouse cells and tissues, bevacizumab can be used to evoke antigen-independent hIgG1 activities.^11^ Consistent with this model, bevacizumab treatment inhibited mouse Vegf-a-induced chemotaxis in bone marrow derived macrophages (BMDMs) from male wild type (C57BL/6J) mice (**Figure 1a**) but did not inhibit chemotaxis of BMDMs from male *Fcgr1*^−/−^ mice (**Figure 1b**). Compared to male wild-type cells, cells isolated from female C57BL/6J mice exhibited significantly less chemotaxis inhibition following hIgG1 treatment (**Figure 1a**). Female *Fcgr1*^−/−^ cells were unaffected by hIgG1 treatment (**Figure 1b**). hIgG1 also exhibited a more potent inhibitory effect on male compared to female-derived cells in primary BMDMs isolated from BALB/c mice (**Figure 1c**), suggesting the sex difference is not exclusive to the C57BL/6J mouse strain.

**Figure 1.**
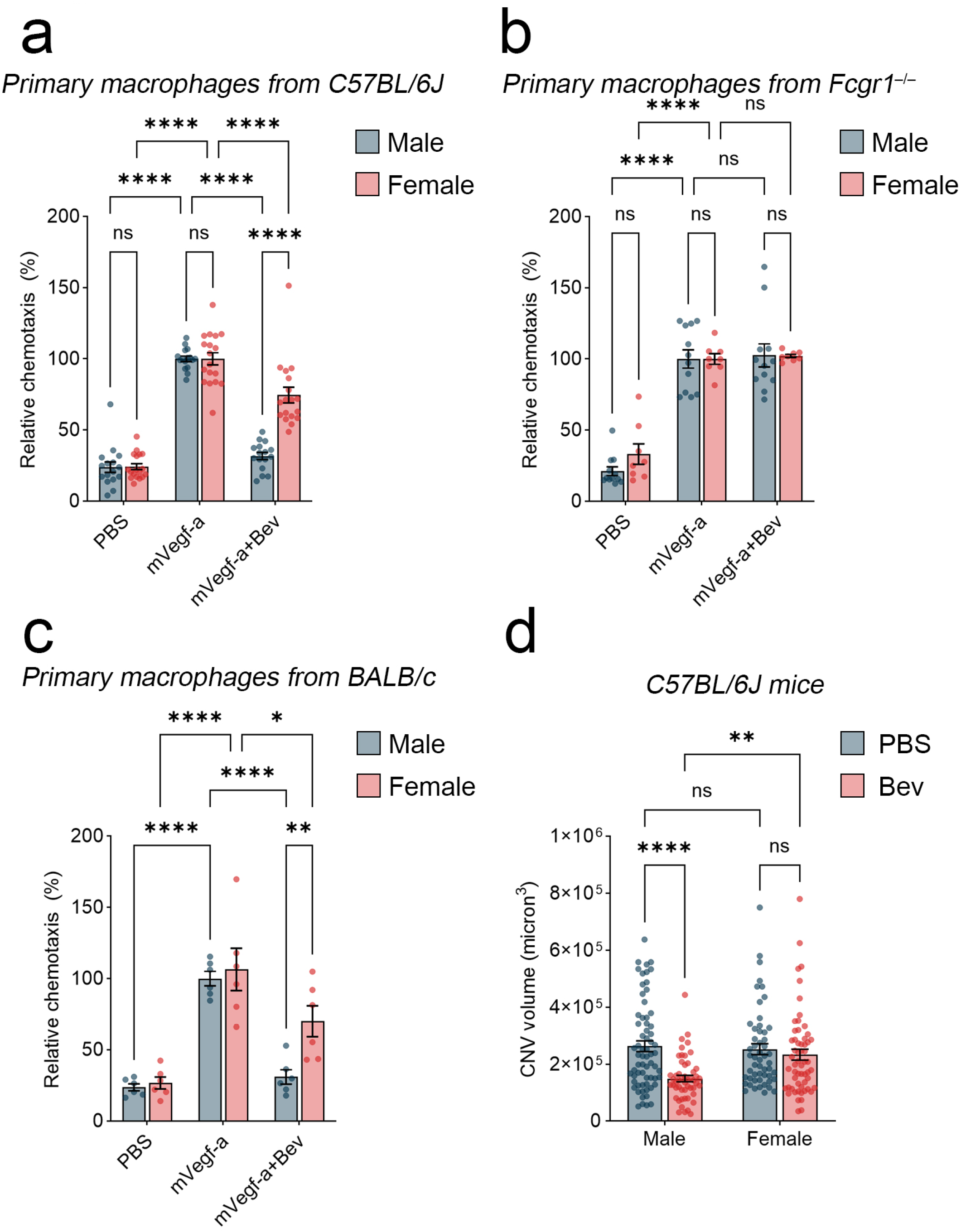
Sex differences in anti-angiogenic activity of hIgG1 in mice. **A**) Relative chemotaxis of bone marrow derived macrophages (BMDM) isolated from C57BL/6J mice from indicated sexes. Chemotaxis was driven by mouse Vegf-a in the presence or absence of the hIgG1 bevacizumab (Bev). Results formulated relative to number of migrated cells after Vegf-a treatment. **B**) Chemotaxis of BMDM isolated from *Fcgr1*^−/−^ mice from indicated sexes. **C**) Chemotaxis of BMDM isolated from BALB/c mice from indicated sexes. **D**) Laser-induced CNV volumes from C57BL/6J mice of indicated sex with intravitreous injection of PBS or Bev. ns, not significant, *P<0.05, **P<0.01, ****P<0.0001.

Recruitment of angiogenic macrophages is essential for neovascular lesion development in choroidal neovascularization (CNV).^17^ Therefore, we hypothesized that hIgG1 would exhibit greater anti-angiogenic effect in male than in female mice. Indeed, intraocular bevacizumab treatment resulted in significantly reduced CNV volume in male, but not female C57BL/6J mice (**Figure 1d**).

Collectively, these findings indicate that hIgG1/ FcγR1-induced chemotaxis and angiogenesis inhibition exhibits a robust sex difference in mice, where males are more susceptible to hIgG1-induced anti-angiogenic responses.

### Sex differences in anti-angiogenic hIgG1 responses in human cells

Antigen-independent hIgG1 suppression of angiogenic processes is also observed in human peripheral blood mononuclear cell (PBMC)-derived macrophages.^11^ These prior studies investigated pooled human PBMCs, isolated what was presumed to be a mixture of male and female donors, and immortalized THP-1 cells which were originally isolated from a male. Thus, to test whether the sex differences identified in mouse BMDM and CNV models was also present in primary human macrophages, we evaluated the effect of hIgG1 on single donor primary CD14^+^, CD16^−^ PBMCs, from males or females, that were differentiated into macrophages by M-CSF. Instead of bevacizumab, which potently neutralizes human VEGF-A,^22^ hIgG1 purified from plasma of a donor with myeloma (Athens Research and Technology) was used. In contrast to hIgG1, hIgG2 does not bind to FcγR1,^23^ and does not inhibit angiogenesis.^11^ Therefore we also treated cells with hIgG2, purified from plasma of a different donor with myeloma, as a negative control. The effect of donor sex and treatment on macrophage chemotaxis are displayed in **Figure 2a**. Consistent with the sex differences observed in mouse models, hIgG1-induced chemotaxis inhibition was significantly greater in macrophages from male compared to female donors (p<0.001). hIgG2 treatment had a comparatively modest effect on chemotaxis and had no sex bias. Similarly, hIgG1 suppressed chemotaxis towards a VEGF-A gradient to a greater degree in macrophages differentiated from THP-1, immortalized from a male donor, compared to HL-60, immortalized from a female donor (**Figure 2b**). hIgG2 treatment did not affect migration in THP-1 cells (**Figure 2c**). Together, these data support that male-specific effects of hIgG1 are conserved in mice and human cells.

**Figure 2.**
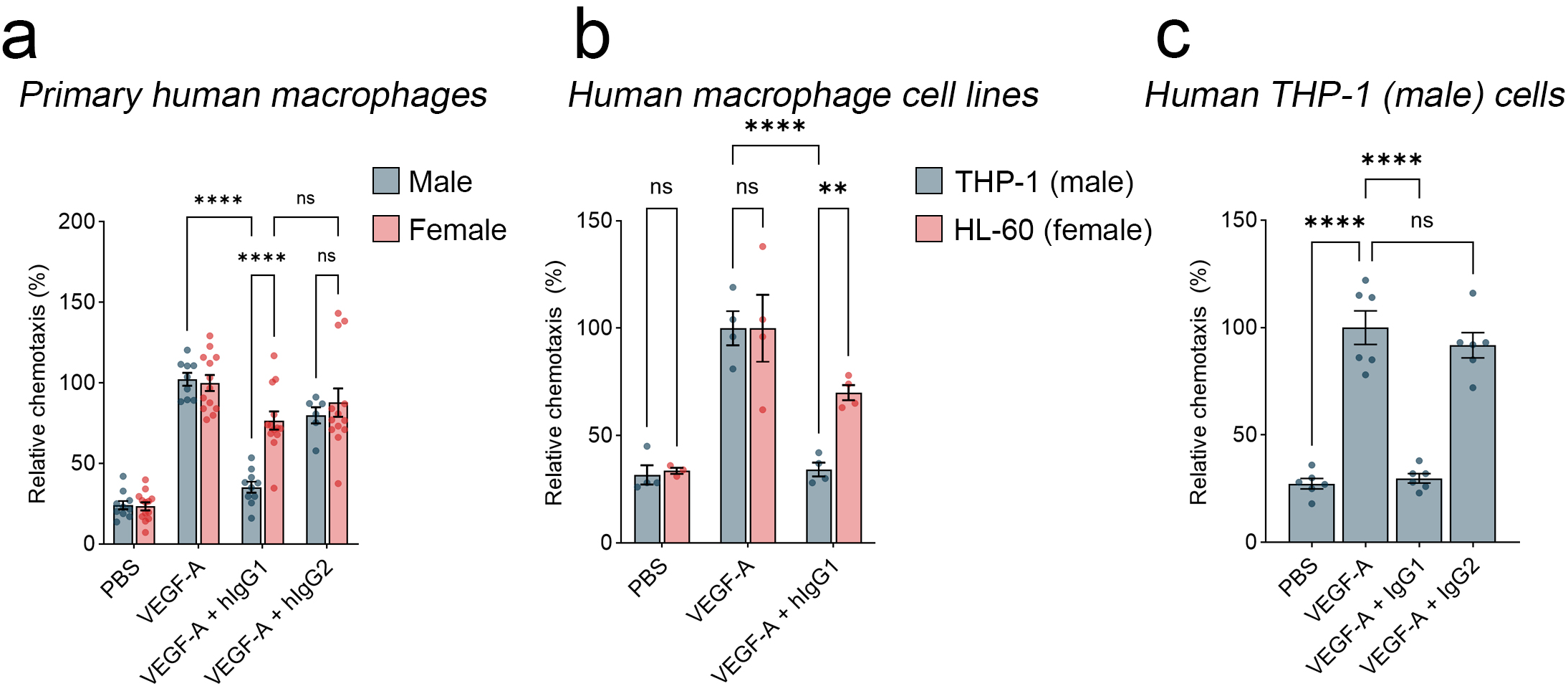
Sex differences in anti-angiogenic activity of hIgG1 in human macrophages. **A)** Relative chemotaxis of primary CD14^+^ macrophages isolated from healthy human donors of indicated sexes. Chemotaxis was driven by human VEGF-A in the presence or absence of hIgG1 or hIgG2 purified from myeloma donors. **B**) Relative chemotaxis of macrophages from THP-1 (male) or HL-60 (female) cell lines driven by human VEGF-A in the presence or absence of hIgG1. **C**) THP-1 macrophage chemotaxis driven by human VEGF-A in the presence or absence of hIgG1 or hIgG2. ns, not significant, **P<0.01, ****P<0.0001.

### Sexual dimorphism in hIgG1 angiogenesis inhibition is mediated in part by the Y chromosome

We next sought to identify the biasing factors responsible for these sex differences in hIgG1-induced angiogenesis inhibition by evaluating this process in the four core genotypes (FCG) model system, which can distinguish between the effects of gonadal hormone secretions and sex chromosome complement on sexually dimorphic phenotypes.^13,24^ We performed laser-induced CNV in FCG mice and analyzed the effect of bevacizumab treatment on angiogenesis. As in wild-type mice, bevacizumab significantly reduced CNV volume in XYM (mice with XY chromosomes and testes) but not XXF (mice with XX chromosomes and ovaries) (**Figure 3a**). Bevacizumab did not significantly reduce CNV volume in either XYF or XXM mice, implying interacting contributions of sex chromosome complement and gonadal secretions to the sex differences in hIgG1-mediated angiogenesis inhibition.

**Figure 3.**
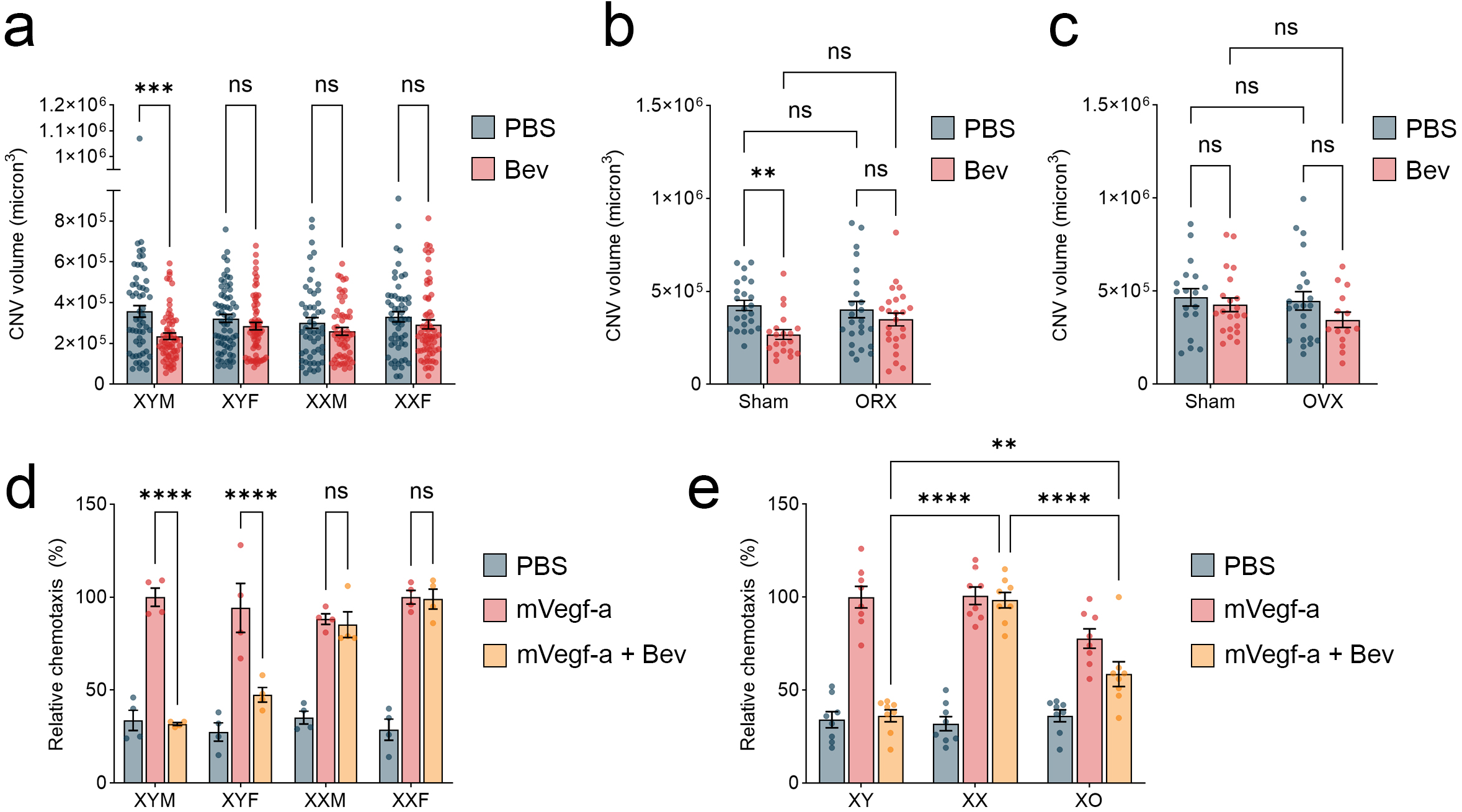
Sex chromosome complement contributes to sex differences in hIgG1 responses. **A**) Laser-induced CNV volume in four core genotypes mice of indicated sex chromosome complement and sex organs (M = testes/penis, F = ovaries) with intravitreous injection of PBS or Bev. **B**) Laser-induced CNV volume in male C57BL/6J mice following orchiectomy or sham surgery with intravitreous injection of PBS or Bev. **C**) Laser-induced CNV volume in female C57BL/6J mice following ovariectomy or sham surgery with intravitreous injection of PBS or Bev. **D**) Relative chemotaxis of BMDM isolated from four core genotypes mice of indicated sex chromosome complement and sex organs. Chemotaxis was driven by mouse Vegf-a in the presence or absence of the hIgG1 bevacizumab (Bev). **E**) Relative chemotaxis of BMDM isolated from XY* Turner syndrome mice of indicated sex chromosome complement. (XO = one single X chromosome). Chemotaxis was driven by mouse Vegf-a in the presence or absence of the hIgG1 bevacizumab (Bev). ns, not significant, **P<0.01, ***P<0.001, ****P<0.0001.

Gonadal secretions can elicit both permanent and reversible sex-biased traits. To test whether reversible effects of gonadal secretions contribute to the sex difference in hIgG1-mediated angiogenesis inhibition, we assessed whether gonadectomy affected bevacizumab-induced CNV suppression. Neither orchiectomy nor ovariectomy affected baseline CNV in young male and female mice respectively (**Figure 3b-c**). Bevacizumab’s anti-angiogenic activity was impaired by orchiectomy in male mice (**Figure 3b**) but was not changed by ovariectomy in female mice (**Figure 3c**). These findings suggest that interactions between irreversible genetic and/or reversible gonadal secretion-dependent differences contribute to sex differences between males and females in hIgG1 responses.

To identify mechanisms responsible for irreversible sex differences in hIgG1 responses, we next quantified hIgG1-induced chemotaxis inhibition in macrophages isolated from the FCG mice. As in laser CNV, macrophages from XYM were susceptible and macrophages from XXF were resistant to bevacizumab-induced chemotaxis inhibition (**Figure 3d**). Interestingly, while XXM macrophages were resistant, XYF macrophages were susceptible to bevacizumab-induced chemotaxis suppression. These findings indicate that differences in sex chromosome complement (XX vs. XY) determines susceptibility to hIgG1-induced chemotaxis inhibition.

Sex differences that depend on chromosome complement may primarily arise due to a) the presence versus absence of a Y chromosome in males and females respectively, or b) the presence of two versus one X chromosomes in females and males respectively. To clarify which scenario is responsible for sex differences in hIgG1 chemotaxis inhibition, we isolated macrophages from mice with X monosomy (XO) produced from the XY* (Turner syndrome) mouse model,^24^ and compared BMDM isolated from wild-type male (XY), female (XX), and female XO mice, which have neither a Y chromosome nor a second X chromosome. Like macrophages from XX mice, XO cells were resistant to bevacizumab-induced chemotaxis suppression (**Figure 3e**). Because the chemotaxis inhibition responses in XO cells were significantly diminished compared to XY cells, we interpreted this to mean that the presence of the Y chromosome contributes to the skewed susceptibility of male cells to hIgG1 chemotaxis inhibition.

### Sexual dimorphism in hIgG1 chemotaxis inhibition depends on Y chromosome-encoded Ddx3y

To identify individual Y chromosome-encoded genes contributing to hIgG1 responses, we performed a transcriptomic screen of the mouse Y chromosome, which contains just 14 known protein-coding genes,^25^ each of which, except for *Eif2s3y*, has a human Y chromosome-encoded homolog. Using qRT-PCR of wild-type male mouse BMDM, we detected expression of 8/13 genes (**Figure 4a**). Among these, we performed an initial siRNA-based screen of four genes (*Ddx3y, Kdm5d, Rbm31y*, and *Uty*), while 4 other genes (*Zfy1, Ssty1, Ssty2*, and *Usp9y*) were poor candidates for reliable siRNA design due to excess homology to other genes or internally repetitive sequences. Using human THP-1 macrophages and primary mouse BMDM, we identified DDX3Y/Ddx3y and UTY/Uty targeted siRNAs as able to reliably abrogate hIgG1 responses (**Figure 4b**). Consistent with reported transcriptional interdependence of the Uty/Ddx3y locus,^26^ silencing *Uty* by either of two siRNAs also suppressed *Ddx3y* abundance. However, *Ddx3y* silencing did not suppress *Uty* (**Figure 4c**). Therefore, without excluding a role for UTY/Uty in this process, we chose to focus on DDX3Y/Ddx3y as a mediator of hIgG1-induced angiogenesis inhibition.

**Figure 4.**
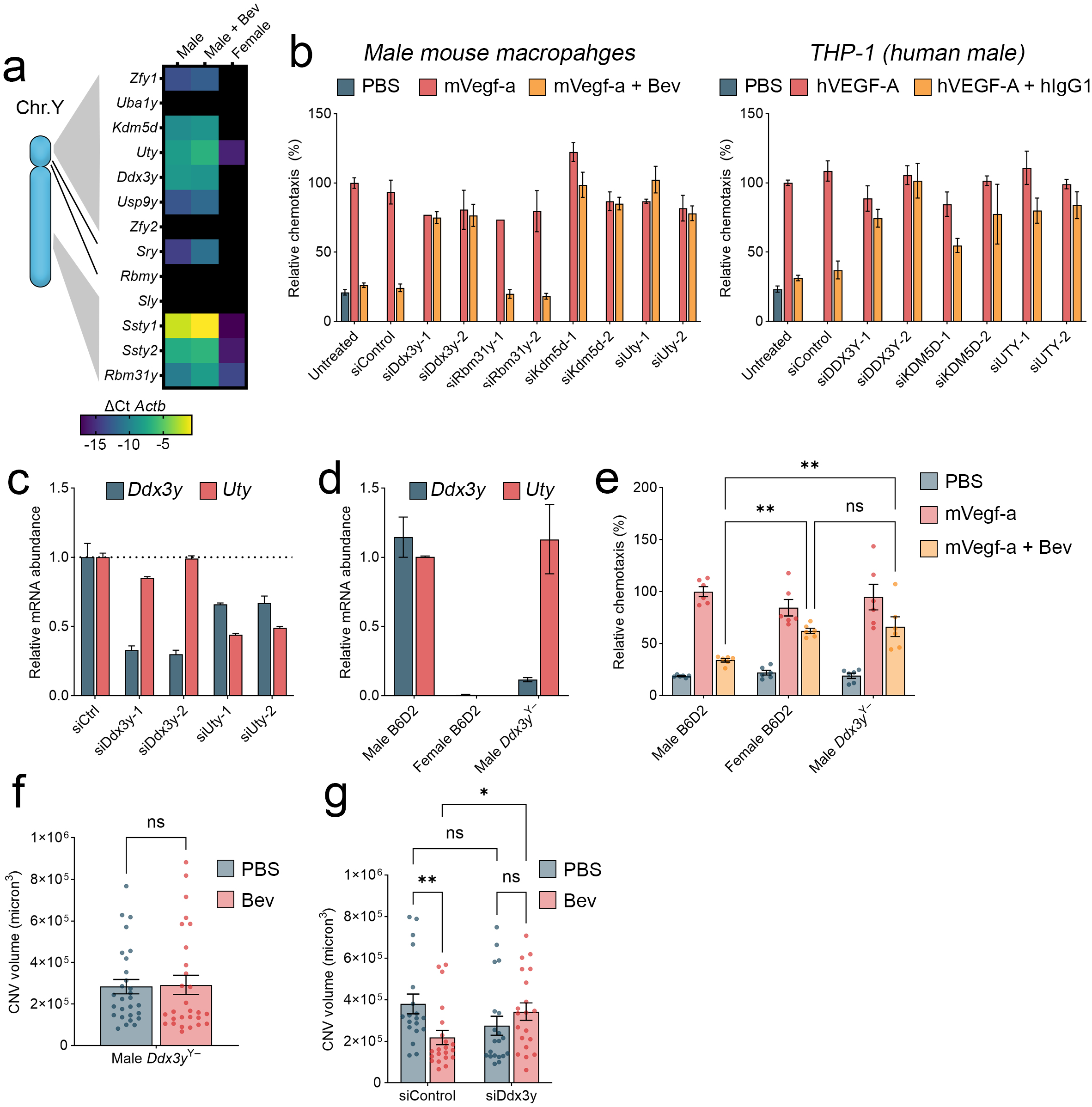
DDX3Y contributes to sex differences in hIgG1 responses. **A**) Transcriptomic screen of 13 mouse Y chromosome protein coding genes. Relative mRNA abundance was quantified by qRT-PCR relative to the housekeeping gene *Bact* in BMDM from male and female mice with or without treatment of bevacizumab (Bev). **B**) siRNA screening of Y chromosome encoded genes. Primary male mouse BMDM (left) or human THP-1 (right) were transfected with two independent siRNAs targeting indicated Y chromosome genes or a scrambled control (siControl). Chemotaxis was assessed in response to mouse Vegf-a or human VEGF-A and bevacizumab (Bev) or hIgG1 as indicated. **C**) mRNA abundance of *Ddx3y* and *Uty* after transfection with siRNAs targeting Ddx3y or Uty, or a scrambled control (siCtrl). **D**) mRNA abundance of *Ddx3y* and *Uty* in BMDM isolated from male wild-type (B6/D2), female B6/D2, or Ddx3y knockout mice. **E**) Relative chemotaxis of BMDM isolated from male B6/D2, female B6/D2, or Ddx3y knockout mice. Chemotaxis was driven by mouse Vegf-a in the presence or absence of Bev. **F**) Laser-induced CNV in Ddx3y knockout mice with intravitreous injection of PBS or Bev. **G**) Laser-induced CNV in wild-type mice with intravitreous injection of cell permeable siRNA targeting Ddx3y (vs. non-targeting control) with PBS or Bev. ns, not significant, *P<0.05, **P<0.01.

To validate findings from the siRNA screen, we next isolated macrophages from a *Ddx3y*^Y–^ mouse line, originally created by CRISPR-mediated targeting of exon 4 of Ddx3y,^27^ as well as female littermates and wild-type B6/D2 *Ddx3y*^Y+^ male mice. *Ddx3y*^Y–^ exhibited a ∼10-fold reduction in *Ddx3y* mRNA compared to *Ddx3y*^Y+^ male BMDM by qRT-PCR (**Figure 4d**). Similar to *Ddx3y* silencing by siRNA, *Ddx3y* genetic ablation did not affect *Uty* mRNA abundance compared to B6/D2 *Ddx3y*^Y+^ male macrophages (**Figure 4d**). Bevacizumab inhibited chemotaxis in BMDM from B6/D2 *Ddx3y*^Y+^ male mice, but *Ddx3y*^Y-^ and female macrophages were relatively resistant (**Figure 4e**). Male *Ddx3y*^Y–^ mice were resistant to the inhibitory effect of bevacizumab on laser-induced CNV (**Figure 4f**). Similarly, intravitreous administration of a cell-permeable cholesterol-conjugated *Ddx3y-*targeted siRNA abrogated bevacizumab responses in wild-type C57BL/6J male mice (**Figure 4g**). Transcripts of the related mouse-specific paralogous gene *D1pas1*, encoded on autosomal chromosome 1 and which may possess overlapping functions in spermatogenesis,^27^ were undetectable (Ct>36) in male BMDM.

DDX3Y silencing by either of two independent siRNAs also reduced hIgG1-induced chemotaxis suppression in primary macrophages derived from healthy male donors (**Figure 5a**). Therefore, we identified DDX3Y silencing as sufficient to abrogate hIgG1-induced chemotaxis suppression.

**Figure 5.**
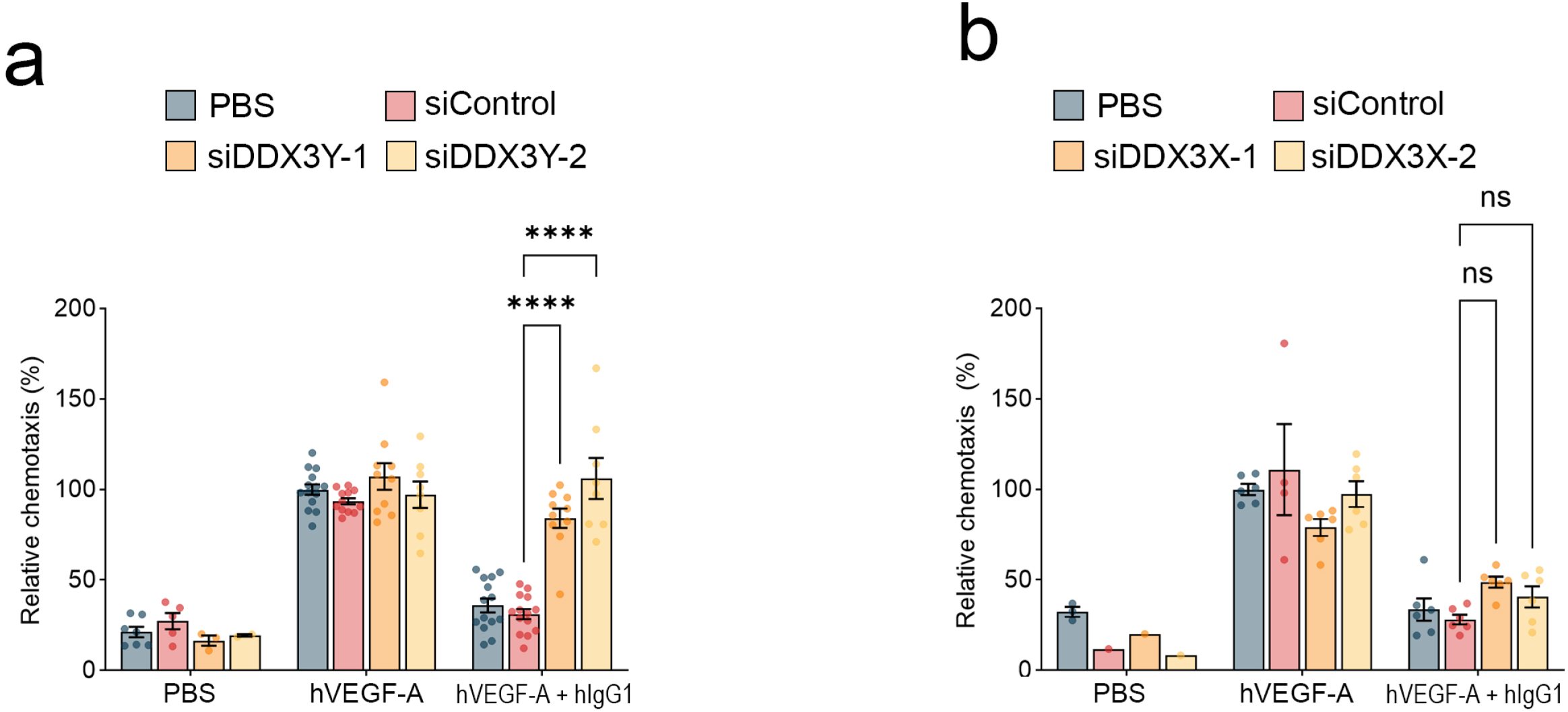
DDX3Y contributes to sex differences in hIgG1 responses in human macrophages. **A**) Relative chemotaxis of primary human macrophages isolated from healthy male donors following transfection with siRNAs targeting DDX3Y or scrambled control (siControl). Chemotaxis was driven by human VEGF-A in the presence or absence of hIgG1. **B**) Relative chemotaxis of male THP-1 macrophages following transfection with siRNAs targeting the X chromosome-encoded DDX3X or scrambled control (siControl). Chemotaxis was driven by human VEGF-A in the presence or absence of hIgG1. ns, not significant, ****P<0.0001.

DDX3Y is a male-specific paralog of DDX3X, a gene that is critical for macrophage hematopoiesis and innate immune responses.^28,29^ siRNAs targeting DDX3X did not impair male THP-1 macrophage responses (**Figure 5b**), supporting the specificity of these reagents. Together, these findings implicate Y chromosome-encoded Ddx3y as contributing to sex differences in hIgG1-induced macrophage chemotaxis and angiogenesis suppression.

## Discussion

We report that the anti-angiogenic activity of hIgG1 antibodies is greater in males than in females, and this activity depends on the Y chromosome. These studies add to the growing literature that Y chromosome genes possess functional activities in non-reproductive tissues. For example, mosaic loss of chromosome Y (mLOY), the most common somatic mutation in men, is age- and smoking-related, and is an independent risk factor for age-related macular degeneration, Alzheimer’s disease, cardiac fibrosis, and premature death.^30^

Our findings revealed that the Y chromosome-encoded gene DDX3Y conferred the inhibitory effects of hIgG1 on angiogenesis. The contribution of individual Y chromosome genes for disease processes is emerging. For example, DDX3Y has been associated with B cell lymphoma,^31,32^ neuronal differentiation,^33^ and graft versus host disease.^34^ Similarly, reduced abundance of the neighboring gene UTY is implicated in heart failure.^35^ Much of DDX3Y function has been inferred from its paralog X chromosome encoded DDX3X, with which it shares 92% amino acid homology. Evidence suggests that these proteins exhibit functional redundance, as DDX3Y can partially compensate for loss of DDX3X. DDX3 RNA helicases have established functional roles in as transcriptional regulators and translation-initiation factors.^32,36^ Still, differences in their N terminal intrinsically disordered regions leads to greater stress granule stability and translation repression activity in DDX3Y compared to DDX3X, which may contribute to sex differences in RNA metabolism.^37,38^ The extent to which DDX3Y contributes to hIgG1-mediated angiosupression via its translated protein remains a difficult question to study given the intrinsic challenges associated with protein-based analyses of Y chromosome encoded genes.^39,40^

Sex differences have been identified in the prevalence, severity, prognosis, and even preferred treatment options for many complex diseases such as peripheral artery disease (PAD),^41^ neovascular AMD,^42^ various cancers,^43^ coronary artery disease,^44,45^ stroke,^46^ cerebral palsy,^47^ and heart failure.^48^ Similarly, a wide range of sex-specific phenotypes have been reported for diverse angiogenic conditions in animal models, including greater vascularity of lung tumors in females,^49^ impaired angiogenesis in males during hyperoxic lung injury,^50–52^ accelerated corneal neovascularization in males after chemical cauterization,^53^ and greater vascularity of perigonadal adipose in females fed a high-fat diet.^54^ Our study suggests that DDX3Y-dependent angiogenesis inhibition may be a sex biasing factor affecting immune-vascular trafficking, and is therefore a candidate to contribute to sex differences in such conditions. Together, our findings suggest that greater female susceptibility to neovascular AMD may arise due to reduced angioregulatory activity of hIgG1 antibodies, which is mediated in part by the Y chromosome gene DDX3Y. Because macrophage chemotaxis is critical in a variety of neovascular settings, this activity may also contribute to sex differences more broadly in diseases of angiogenesis.

## Notes

### Competing Interest Statement

J.A. is a co-founder of iVeena Holdings, iVeena Delivery Systems and Inflammasome Therapeutics and, unrelated to this work, has been a board member for Theragen and consultant for Abbvie/Allergan, Retinal Solutions and Saksin LifeSciences.

